# Senolytic therapy alleviates physiological human brain aging and COVID-19 neuropathology

**DOI:** 10.1101/2023.01.17.524329

**Authors:** Julio Aguado, Alberto A. Amarilla, Atefeh Taherian Fard, Eduardo A. Albornoz, Alexander Tyshkovskiy, Marius Schwabenland, Harman K. Chaggar, Naphak Modhiran, Cecilia Gómez-Inclán, Ibrahim Javed, Alireza A. Baradar, Benjamin Liang, Malindrie Dharmaratne, Giovanni Pietrogrande, Pranesh Padmanabhan, Morgan E. Freney, Rhys Parry, Julian D.J. Sng, Ariel Isaacs, Alexander A. Khromykh, Alejandro Rojas-Fernandez, Thomas P. Davis, Marco Prinz, Bertram Bengsch, Vadim N. Gladyshev, Trent M. Woodruff, Jessica C. Mar, Daniel Watterson, Ernst J. Wolvetang

## Abstract

Aging is the primary risk factor for most neurodegenerative diseases, and recently coronavirus disease 2019 (COVID-19) has been associated with severe neurological manifestations that can eventually impact neurodegenerative conditions in the long-term. The progressive accumulation of senescent cells *in vivo* strongly contributes to brain aging and neurodegenerative co-morbidities but the impact of virus-induced senescence in the aetiology of neuropathologies is unknown. Here, we show that senescent cells accumulate in physiologically aged brain organoids of human origin and that senolytic treatment reduces inflammation and cellular senescence; for which we found that combined treatment with the senolytic drugs dasatinib and quercetin rejuvenates transcriptomic human brain aging clocks. We further interrogated brain frontal cortex regions in postmortem patients who succumbed to severe COVID-19 and observed increased accumulation of senescent cells as compared to age-matched control brains from non-COVID-affected individuals. Moreover, we show that exposure of human brain organoids to SARS-CoV-2 evoked cellular senescence, and that spatial transcriptomic sequencing of virus-induced senescent cells identified a unique SARS-CoV-2 variant-specific inflammatory signature that is different from endogenous naturally-emerging senescent cells. Importantly, following SARS-CoV-2 infection of human brain organoids, treatment with senolytics blocked viral retention and prevented the emergence of senescent corticothalamic and GABAergic neurons. Furthermore, we demonstrate in human ACE2 overexpressing mice that senolytic treatment ameliorates COVID-19 brain pathology following infection with SARS-CoV-2. *In vivo* treatment with senolytics improved SARS-CoV-2 clinical phenotype and survival, alleviated brain senescence and reactive astrogliosis, promoted survival of dopaminergic neurons, and reduced viral and senescence-associated secretory phenotype gene expression in the brain. Collectively, our findings demonstrate SARS-CoV-2 can trigger cellular senescence in the brain, and that senolytic therapy mitigates senescence-driven brain aging and multiple neuropathological sequelae caused by neurotropic viruses, including SARS-CoV-2.

## Introduction

Although severe acute respiratory syndrome coronavirus 2 (SARS-CoV-2) is primarily a respiratory viral pathogen and the cause of coronavirus disease 2019 (COVID-19), persistent post-acute infection syndromes (PASC) derived from viral infections including SARS-CoV-2 are emerging as a frequent clinical picture^1,2^. In fact, most COVID-19 patients including individuals with or without comorbidities, and even asymptomatic patients, often experience a range of neurological complications^3,4^. ‘Long-COVID’ is a type of PASC that is gaining significant awareness, with patients reporting persistent manifestations, such as hyposmia, hypogeusia, sleep disorders and substantial cognitive impairment, the latter affecting approximately one in four COVID-19 cases^5–7^. These clinical symptoms are supported by ample evidence of SARS-CoV-2 infectivity in multiple cell types of the nervous system^8–16^ and significant structural changes in the brains of COVID-19 patients^17^. Furthermore, patient transcriptomic data from *postmortem* brain tissue indicate associations between the cognitive decline observed in patients with severe COVID-19 and molecular signatures of brain aging^18^. In agreement with this observation, *postmortem* patient biopsies show that SARS-CoV-2-infected lungs — compared to uninfected counterparts — accumulate markedly higher levels of senescence^19^; a cellular phenotype known to contribute to organismal aging^20^ and co-morbidities such as chronic degenerative conditions^21^. Importantly, although recent data supports a role for senescent cells in driving neurodegeneration and cognitive decline in *in vivo* models of neuropathology^22,23^ and in physiologically aged mice^24^, their contribution to COVID pathology in the central nervous system (CNS) and human tissue brain aging remains unknown.

In the past decade, numerous strategies have been developed to target senescent cells^25^. Among these, the pharmacological removal of senescent cells with senolytic drugs has become one of the most explored interventions, with many currently in human clinical trials^26^. A group of these senolytics — such as the cocktail of dasatinib plus quercetin (D+Q), or fisetin — exhibit blood-brain barrier permeability upon oral administration^22,27^, making these formulations particularly valuable to test the contribution of senescence in the brain *in vivo*.

In the present study, we first document the efficacy of multiple senolytic interventions in clearing senescent cells in physiologically aged human pluripotent stem cell-derived brain organoids. Transcriptomic analysis across individual senolytic treatments revealed a differential effect in modulating the senescence-associated secretory phenotype (SASP), with a distinctive impact of D+Q administration in rejuvenating the organoids transcriptomic aging clock. Importantly, we report an enrichment of senescent cells in *postmortem* brain tissue of COVID-19 patients and further show a direct role for SARS-CoV-2 and highly neurotropic viruses such as Zika and Japanese encephalitis in evoking cellular senescence in human brain organoids. SARS-CoV-2 variant screening identified Delta (B.1.617.2) as the variant that exerts the strongest induction of cellular senescence in human brain organoids, and spatial transcriptomic analysis of Delta-induced senescent cells unveiled a novel type of senescence that exhibits a different transcriptional signature from senescent cells that naturally emerge in *in vitro* aged uninfected organoids. Furthermore, senolytic treatment of SARS-CoV-2-infected organoids selectively removed senescent cells, lessened SASP-related inflammation and reduced SARS-CoV-2 RNA expression, indicating a putative role for senescent cells in facilitating viral retention. Finally, to gain *in vivo* relevance of these findings, we examined the treatment effects of senolytics in transgenic mice expressing human angiotensin-converting enzyme 2 (hACE2)^8^ previously infected with SARS-CoV-2 and observed improved clinical performance and survival, reduced viral load in the brain, improved survival of dopaminergic neurons, decreased astrogliosis, and attenuated senescence and SASP gene expression in the brains of the infected mice. Our findings suggest a detrimental role for virus-induced senescence in accelerating brain inflammation and the aging process in the CNS, and a potential therapeutic role for senolytics in the treatment of COVID-19 neuropathology.

## Results

### Senolytics target biological aging and senescent cells in physiologically aged human brain organoids

To explore the efficacy of senolytics in clearing senescent cells from human brain tissue models, we generated 8-month-old human brain organoids (BOs) from embryonic stem cells and exposed these to two doses of senolytics for one month at 2 weekly intervals (Supplementary Fig. 1a). We tested the Bcl-2 inhibitors navitoclax and ABT-737, as well as D+Q senolytic drug combination, and quantified the abundance of cells exhibiting senescence-associated β-galactosidase activity (SA-β-gal). Exposure to senolytics resulted in significantly lower SA-β-gal activity as compared to vehicle-treated controls (Fig. 1a, c), indicating that all treatments eliminated a large number of senescent cells in the treated BOs. In agreement with this, analysis of lamin B1 protein expression — a nuclear lamina marker often downregulated in senescence^28^ — within organoid sections revealed a significantly higher content of lamin B1 in the senolytic-treated organoids as compared to control counterparts (Fig. 1b, d), further indicating that senolytics cleared senescent cells by enriching for lamin B1^High^ cell populations.

**Figure 1.**
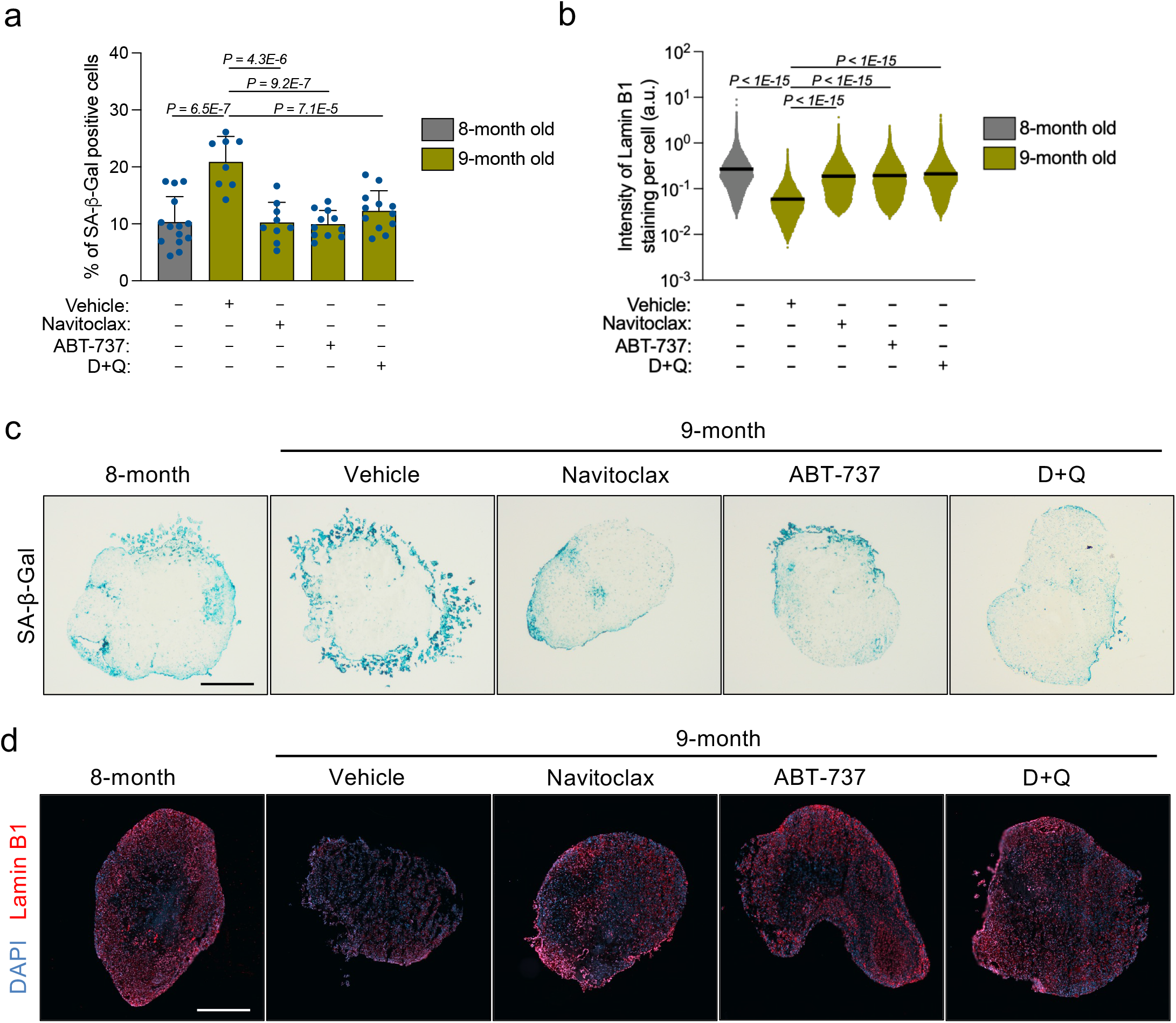
Long-term senolytic treatment prevents selective accumulation of senescent cells in physiologically aged human brain organoids. Brain organoids were generated and grown *in vitro* for 8 months, and subsequently exposed to two doses (each dose every two weeks) of either navitoclax (2.5 μM), ABT-737 (10 μM) or D+Q (D: 10 μM; Q: 25 μM) administration within the following month, after which the organoids were collected for *in situ* analysis. **(a)** SA-β-gal assays were performed on organoid sections. Each data point in the bar graph represents a single organoid analysed. Error bars represent s.d.; at least 8 individual organoids were analysed per condition; one-way ANOVA with Tukey’s multiple-comparison post-hoc corrections. **(b)** Lamin B1 staining was performed on organoid sections. Each data point in the scatter plot represents the integrated intensity of each cell within organoid sections. At least 8 individual organoids were analysed per condition; one-way ANOVA with Tukey’s multiple-comparison post-hoc corrections. **(c,d)** Representative images from quantifications shown in a and b, respectively. Scale bar, 0.3 mm.

We next performed whole-organoid RNA sequencing to compare the transcriptomes of senolytic-treated and vehicle control 9-month-old BOs. Consistent with our protein expression data (Fig. 1b, d), *LMNB1* (lamin B1) mRNA levels were significantly upregulated in all three senolytic-treated organoids compared to vehicle-treated counterparts (Fig. 2a-c). We further identified 81 senescence-associated genes (including the proinflammatory genes *CXCL13* and *TNFAIP8*) that were consistently suppressed upon all three senolytic interventions (Fig. 2d and Supplementary Fig. 1b). We however also noticed that each senolytic treatment exerted substantially different effects in modulating the SASP and other senescence-associated genes (Fig. 2a-c). For instance, *SERPINF1* was significantly repressed upon ABT-737 administration (Fig. 2b) while D+Q did not modulate *SERPINF1* expression but greatly supressed *IL8, SERPINE1* and *IL1A* (Fig. 2c). Compared to navitoclax and ABT-737 – compounds that modulate multiple shared genes that are enriched for a few pathways (e.g. K-Ras signalling) (Fig. 2e) –, D+Q had a broader spectrum effect, mitigating multiple pro-inflammatory pathways characteristic of cellular senescence, such as NF-κB and IFNγ signalling (Fig. 2e and Supplementary Fig. 1c). In addition, we identified mTOR as a significantly supressed pathway upon D+Q treatment (Fig. 2e), validating the effects reported for Q as an inhibitor of mTOR kinase. We next performed aging clock predictions based on whole transcriptome sequencing to further explore the impact of senolytics on the aging process. Remarkably, in addition to their senolytic mechanisms of action, D+Q treatments on 9-month-old organoids reverted their gene expression age to levels comparable of 8-month-old counterparts according to transcriptomic brain aging clock analysis (Fig. 2f), a phenotype not recapitulated by the other two senolytics tested. Besides negative association with aging, gene expression changes induced by D+Q treatment were positively correlated with mammalian signatures of established lifespan-extending interventions, such as caloric restriction and rapamycin administration (Fig. 2g), indicating a health-promoting role of D+Q in targeting cellular senescence and biological aging in human CNS tissues.

**Figure 2.**
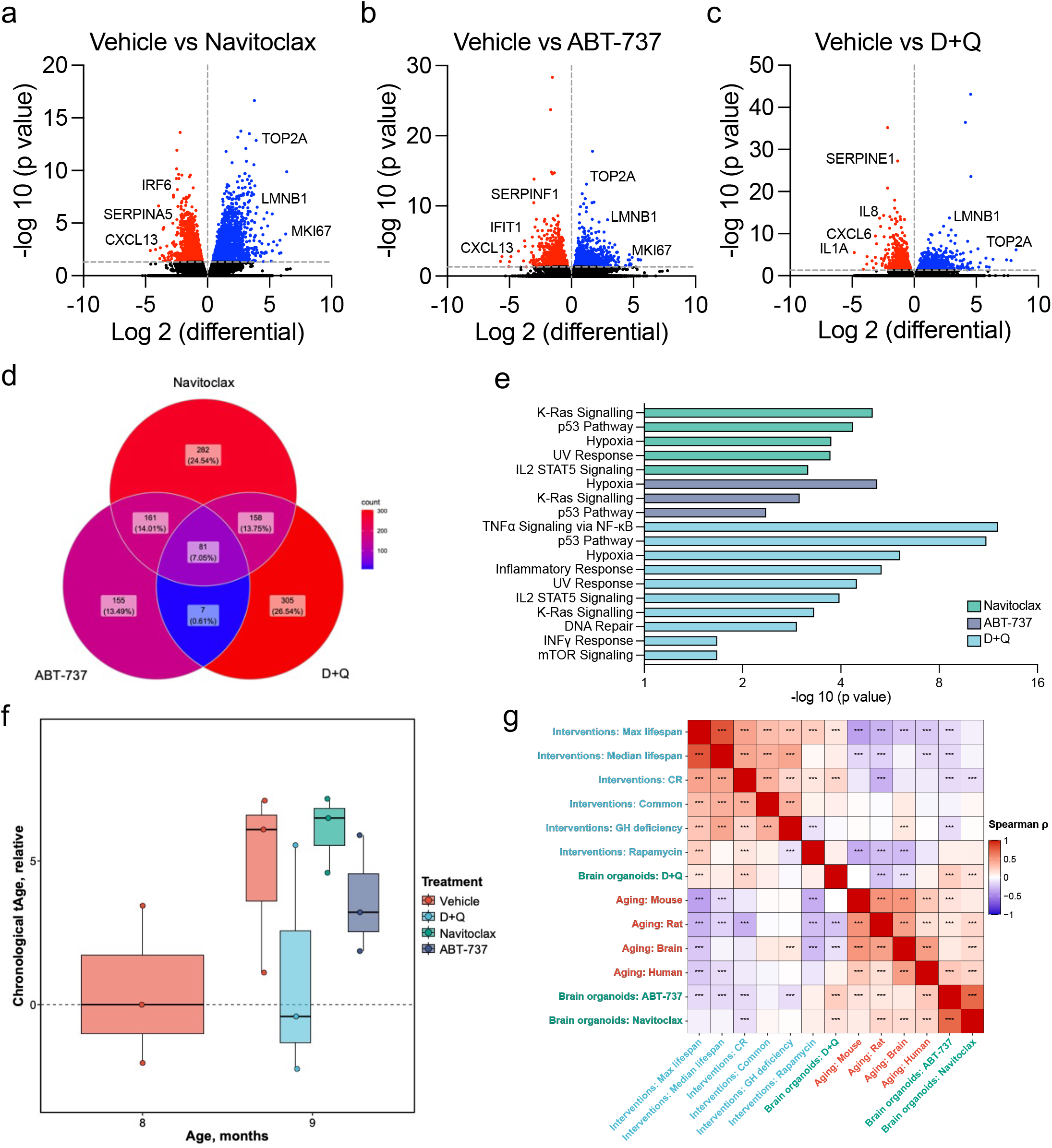
Transcriptomic characterization of distinct senolytic interventions on brain aging hallmarks. Brain organoids were generated and grown *in vitro* for 8 months, and subsequently exposed to two doses (each dose every two weeks) of either navitoclax (2.5 μM), ABT-737 (10 μM) or D+Q (D: 10 μM; Q: 25 μM) administration within the following month, after which the organoids were collected and subjected for bulk RNA sequencing analysis. **(a-c)** Volcano plots show vehicle-treated versus **(a)** navitoclax-, **(b)** ABT-737- and **(c)** D+Q-treated brain organoid differential expression of upregulated (blue) and downregulated (red) genes. **(d)** Venn diagram shows differentially repressed senescence-associated genes among senolytic-treated organoids defined with a significance adjusted P value <0.05. **(e)** Gene Set Enrichment Analysis was carried out using aging hallmark gene sets from the Molecular Signature Database. The statistically significant signatures were selected (FDR <0.25) and placed in order of normalized enrichment score. Bars indicate the pathways enriched in individual senolytic treatments as compared to vehicle-treated brain organoids. **(f)** Transcriptomic age of organoids treated with either vehicle or senolytic compounds assessed using brain multi-species aging clock. **(g)** Spearman correlation between gene expression changes induced by senolytics in aged organoids and signatures of aging and established lifespan-extending interventions based on functional enrichment output. Normalized enrichment scores (NES) calculated with GSEA were used to evaluate correlations between pairs of signatures.

### SARS-CoV-2 infection triggers cellular senescence in the brains of COVID-19 patients and in human brain organoids

Given the observed neuroinflammatory effects of SARS-CoV-2 infection during acute COVID-19 disease^29^ and its association with molecular signatures of aging in patient brains^18^, we postulated that part of this pro-inflammatory aging-promoting environment is brought about by SARS-CoV-2-induced senescence in the brain. To test this hypothesis, we quantified the prevalence of senescent cells in postmortem frontal cortex from age-matched brains of patients that either died following severe COVID-19 or patients who died of non-infectious, and non-neurological reasons. Notably, *in situ* high-throughput analysis of over 2.7 million single cells across 15 individual brain samples (7 COVID-19 and 8 non-COVID-19 frontal cortex sections) revealed increased p16 immunoreactivity frequencies in COVID-19 patient brains, with a >7-fold increase in the number of p16-positive cells as compared to non-COVID-19 age-matched controls (Fig. 3). These results suggest a potential role for SARS-CoV-2 in triggering cellular senescence, a cellular phenotype that contributes to cognitive decline and that could pose a risk in the acceleration of neurodegenerative processes associated with long-COVID.

**Figure 3.**
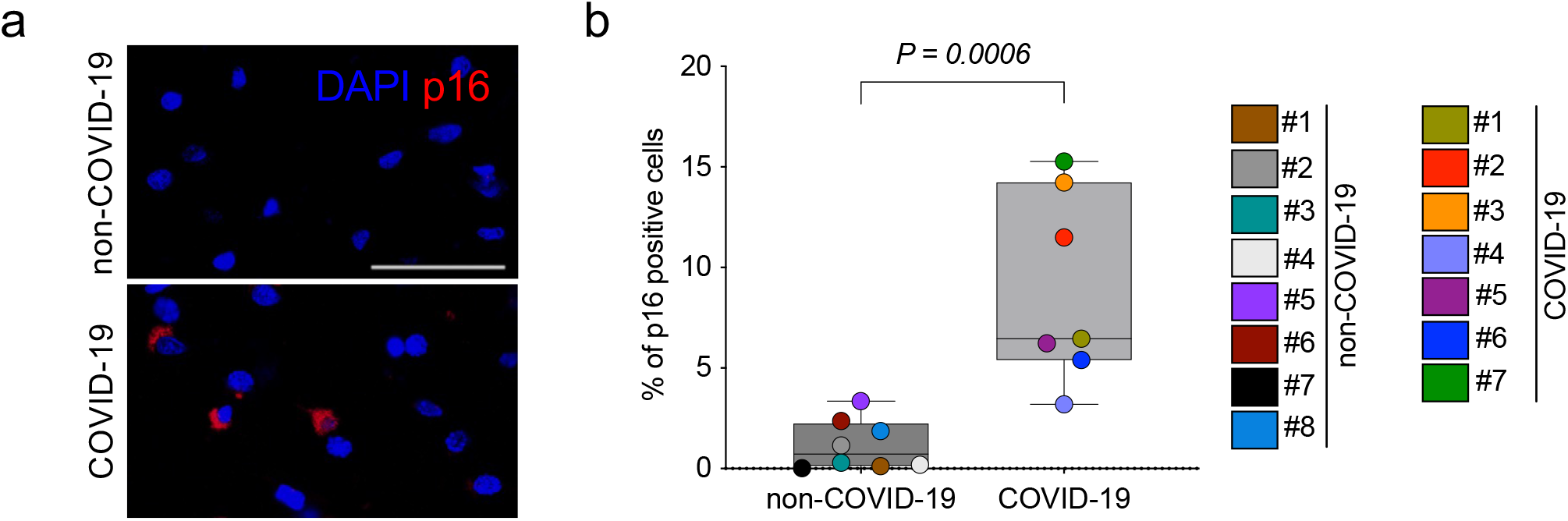
Brains of COVID-19 patients exhibit increased accumulation of p16 senescent cells. **(a)** Immunofluorescence images showing DAPI (blue), and p16 (red) immunoreactivity in sections of frontal cortex regions from patients with severe COVID-19 and age-matched non-COVID-related controls. Scale bar, 50 μm. **(b)** Box plots show the percentage of p16-positive cells. Each data point in the graph represents a single patient analysed, with a total of 2,794,379 individual brain cells across 7 COVID-19 and 8 non-COVID-19 patients. Whiskers represent min-max values; two-tailed Student’s t test.

To mechanistically study the role of neurotropic viruses in aging-driven neuropathology, we exposed human BOs to different viral pathogens, including SARS-CoV-2. Consistent with previous reports^8,9,16,30^, SARS-CoV-2 BO infections were detected largely within populations of neurons and neural progenitors (Supplementary Fig. 2a, b). To test putative virus-induced senescence phenotypes, we screened seven SARS-CoV-2 variants by infecting human BOs at identical multiplicity of infection (MOI) and ranked them based on SA-β-gal activity as initial readouts of cellular senescence. Notably, most variants elicited a significant increase in SA-β-gal, with Delta (B.1.617.2) as the one showing the strongest induction (Fig. 4a, b). In addition, serial sectioning of Delta-infected organoids revealed a distinctive colocalization between SA-β-gal and viral spike protein (Fig. 4c), further supporting a role for SARS-CoV-2 in driving virus-induced senescence in the brain. This phenotype was moreover confirmed when organoid sections were co-immunolabelled with antibodies against p16 and SARS-CoV-2 nucleocapsid antigens (Fig. 4d). Because of the mechanistic role of DNA damage in affecting most aging hallmarks^31^, including the onset of cellular senescence^32^, we next explored whether SARS-CoV-2 infection led to DNA double-strand break accumulation. Consistent with previous evidence^19,33^, we detected significantly heightened levels of phosphorylated histone H2AX at serine 139 (known as γH2AX) in SARS-CoV-2-infected organoid regions as compared to uninfected organoid cells (Fig. 4e, f), indicating increased DNA damage response marks upon SARS-CoV-2 infection. Importantly, virus-induced senescence also became detectable in response to a variety of human neurotropic viruses, including Japanese Encephalitis virus (JEV), Rocio virus (ROCV) and Zika virus (ZIKV) in human BOs (Fig. 4g).

**Figure 4.**
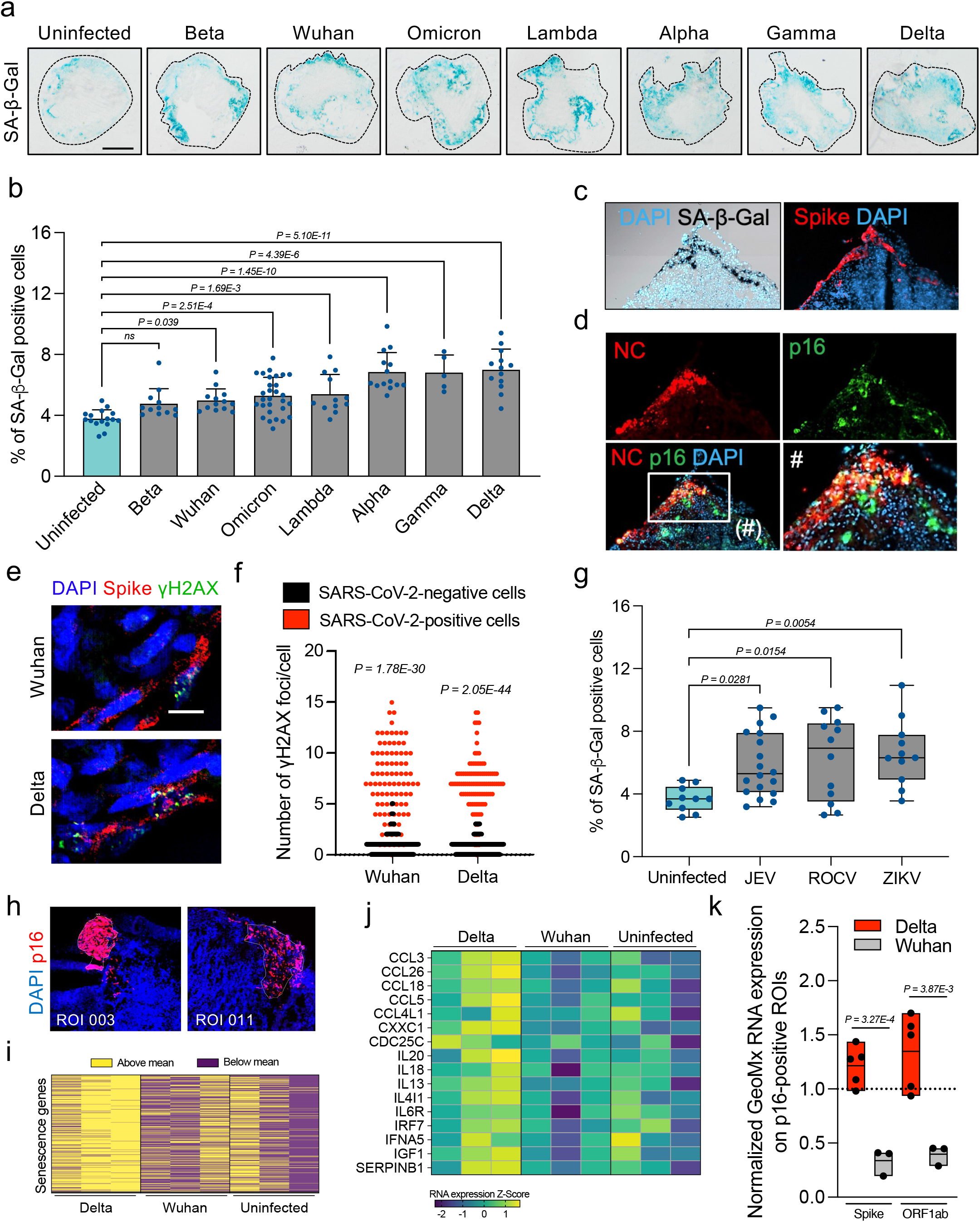
Neurotropic viral infections elicit virus-induced senescence in human brain organoids. **(a)** SARS-CoV-2 variant screening was performed on brain organoids at multiplicity of infection 1 and monitored for SA-β-gal activity at 5 days post infection. Scale bar, 0.3 mm. **(b)** Quantification of data presented in **a.** Bar graphs show the percentage of SA-β-gal-positive cells. Each data point in the bar graph represents a single organoid analysed. Error bars represent s.d.; at least 5 individual organoids were analysed per variant-infected condition; one-way ANOVA with Dunnett’s multiple-comparison post-hoc corrections. **(c)** Representative images of serially sectioned Delta-infected organoid regions stained for SA-β-gal and SARS-CoV-2 spike protein. **(d)** Representative images of the region shown in **c**. co-immunolabelled with p16 and SARS-CoV-2 nucleocapsid (NC) antigen. **(e)** Organoids infected for 5 days with the indicated SARS-CoV-2 variants were stained for γH2AX and SARS-CoV-2 Spike protein. Scale bar, 40 μm. **(f)** Quantification of data presented in **e.** Scatter plot show the number of γH2AX foci per cell in infected regions (red) versus uninfected counterparts (black). Each data point in the scatter plot represents a single cell analysed; at least 400 cells per variant-infected condition have been analysed; two-tailed Student’s t-test. **(g)** Human brain organoids were infected with the neurotropic flaviviruses Japanese Encephalitis virus (JEV), Rocio virus (ROCV) and Zika virus (ZIKV) at multiplicity of infection 0.1; and monitored SA-β-gal activity 5 days post infection. Box plots show the percentage of SA-β-gal-positive cells. Each data point represents a single organoid analysed. Whiskers represent min-max values; at least 5 individual organoids were analysed per virus-infected condition; one-way ANOVA with multiple-comparison post-hoc corrections. **(h-k)** Uninfected, Wuhan- and Delta-infected human brain organoids where subjected to Regions of Interest (ROI) selection based on p16 protein expression for spatial profiling by the Nanostring GeoMX digital spatial profiler assay and further sequenced for the GeoMx Human Whole Transcriptome Atlas. Three organoids were used per condition for ROI selection. **(h)** Representative p16-positive ROIs. **(i)** Heat map of polarity with shown expression above (blue) and below (red) the mean for each differentially heightened SASP gene of Delta-infected p16-positive ROIs. **(j)** Senescence heat map gene expression signature of Delta-infected p16-positive cells. **(k)** Box plots show the expression enrichment of SARS-CoV-2 genes (Spike, ORF1ab) for each SARS-CoV-2 variant. Each data point in the box plot represents a normalized fold change value of SARS-CoV-2 genes on p16-positive ROIs relative to p16-negative counterparts (depicted by a grid line). Whiskers represent min-max values; at least n=3 p16-positive ROIs were analysed per condition; two-tailed Student’s t test.

As SARS-CoV-2 infection is coupled with cognitive decline and signatures of aging, we further assessed associations of transcriptomic changes in COVID-19 patients and SARS-CoV-2-infected human BOs. Specifically, we compared post-mortem frontal cortex transcriptomic data from a COVID-19 cohort study of 44 individual patient brains^18^ with bulk RNA sequencing we performed on human cortical brain organoids 10 days post infection. Notably, among 1,588 differentially expressed genes (DEGs) between SARS-CoV-2-infected human BOs compared and uninfected counterparts, 485 genes (30.54%) were also differentially expressed in COVID-19 patient brain samples. Of note, this common gene set was enriched for known aging and senescence pathways, identified in the hallmark gene set collection of the Molecular Signatures Database^34^ (Supplementary Fig. 3a).

To better understand the differential effects of the ancestral Wuhan virus and Delta (B.1.617.2) SARS-CoV-2 variants on senescence induction in hBOs, performed NanoString GeoMx spatial transcriptomic sequencing on p16 protein-expressing regions of interest (ROIs) within organoid sections (Fig. 4h). ROI selection was performed to enable the capture of targeted transcriptome from sufficient senescent cell tissue (>300 cells per ROI) to generate robust count data. Our bulk RNA sequencing analysis revealed 1,250 DEGs in Wuhan-infected BOs as compared to a lower 474 DEGs in Delta-infected counterparts (Supplementary Fig. 3b), a result possibly explained by the higher infectivity rate observed in the Wuhan-infected organoids (Supplementary Fig. 3c). Strikingly, spatial transcriptome analysis of p16-positive cells identified over 1,100 DEGs in Delta-infected organoids, an effect 100-fold greater than Wuhan where only 9 DEGs were detected (Supplementary Fig. 3b). This was explained by principal component analysis, where gene set space determined that the Delta-infected ROIs were separable from overlapping transcriptomes from Wuhan-infected and uninfected senescent cell regions (Supplementary Fig. 4a). Upon extensive analysis of significantly modulated gene expression in p16-positive ROIs of Delta-infected organoids, we identified 458 genes associated with cellular senescence that differentially clustered from Wuhan-infected and uninfected ROIs (Fig. 4i), with many interleukins significantly elevated in Delta-infected ROIs (Fig. 4j). Importantly, this unique Delta-specific senescence transcriptional signature was detected in the presence of heightened normalized SARS-CoV-2 gene expression in Delta compared to p16-positive cells of Wuhan-infected organoids (Fig. 4k). Altogether, these results demonstrate a direct role for SARS-CoV-2 and neurotropic flaviviruses in fuelling virus-induced senescence, and revealed a specific effect of Delta (B.1.617.2) in inducing the selective induction of a *de novo* transcriptional signature and simultaneous accumulation of SARS-CoV-2 in senescent cells of human BOs.

### Senolytics reduce SARS-CoV-2 viral expression and virus-induced senescence in human brain organoids

The results described so far support a functional role of SARS-CoV-2 in inducing brain cellular senescence. To investigate whether this virus-induced phenotype could be pharmacologically targeted, we next tested the impact of the selective removal of senescent cells with the same senolytic interventions we previously showed were effective in eliminating senescent cells from physiologically aged organoids (Fig. 5a). We observed that senolytic treatments 5 days post SARS-CoV-2 infection significantly reduced the number of brain organoid cells that display SA-β-gal activity (Fig. 5b). Notably, senolytic treatment in Delta-infected organoids had an overall more prominent and statistically significant effect in reducing cellular senescence as compared to Wuhan-infected counterparts, consistent with the stronger virus-induced senescence phenotype observed upon Delta infections in our initial SARS-CoV-2 variant screening (Fig. 5a, b). Moreover, senolytics were able to revert lamin B1 loss induced by Delta infections (Supplementary Fig. 4b). Remarkably, treatment with senolytics reduced the viral load in BOs up to 40-fold as measured by intracellular SARS-CoV-2 RNA levels (Fig. 5c), indicating a putative role of senescent cells as reservoirs that may preferentially facilitate viral replication. To characterise cell type-specific SARS-CoV-2-induced senescence, we performed deconvolution of spatial transcriptomic data from p16-positive cells (Fig. 5d), a type of analysis that enables cell abundance estimates from gene expression patterns^35^. We identified layer 6 corticothalamic neurons (L6CT L6b, > 9-fold induction) and GABAergic ganglionic eminence neurons (CGE, > 4-fold induction) as the two neuronal populations that showed significantly increased senescence incidence upon SARS-CoV-2 infections in brain organoids (Fig. 5e); two brain cell populations that are vital for modulating neural circuitry and processing incoming sensory information^36^. Importantly, all the three senolytic treatments tested prevented the accumulation of cellular senescence in both L6CT L6b and CGE brain organoid cell populations (Fig. 5e).

**Figure 5.**
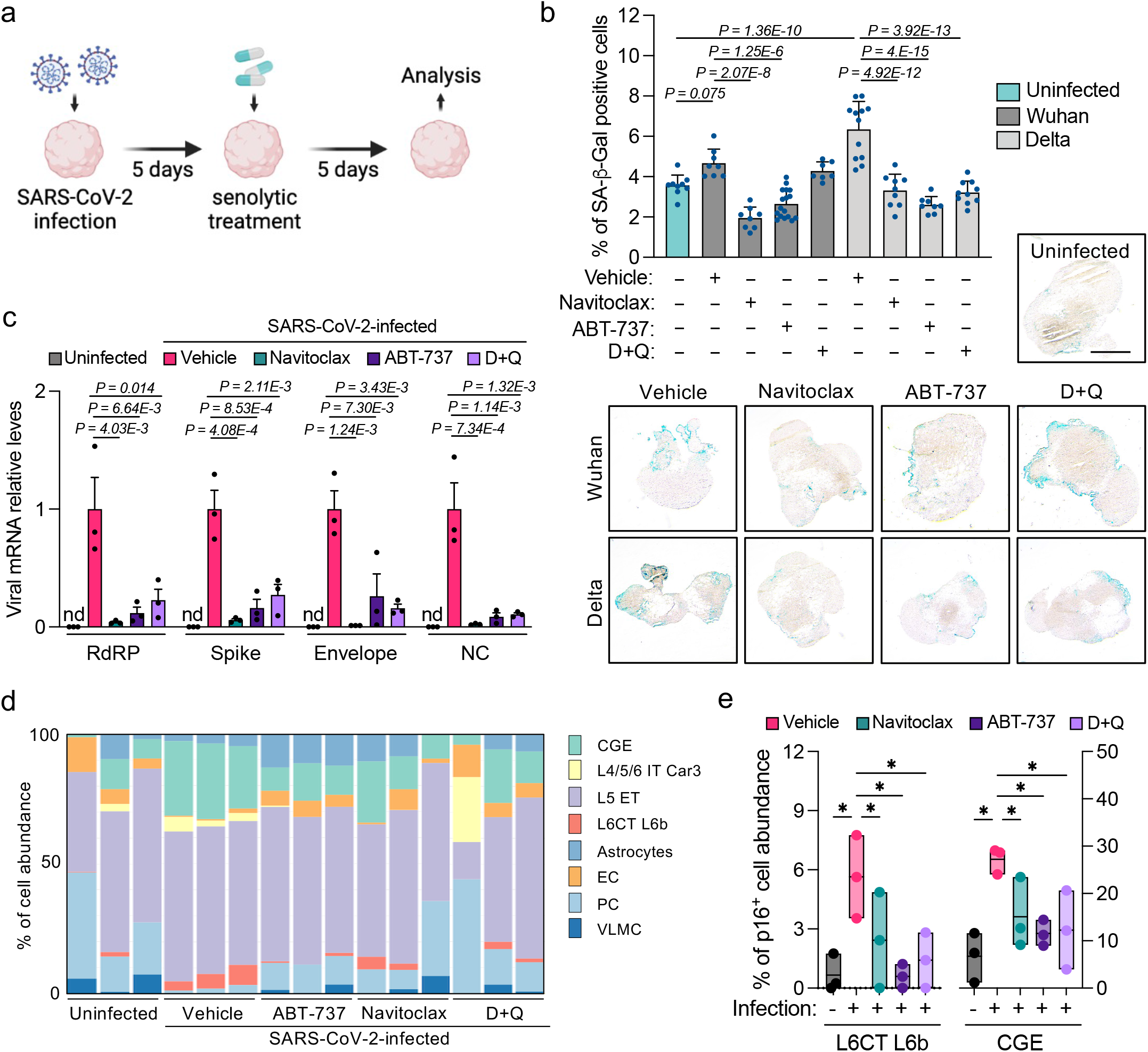
Senolytics clear virus-induced senescence in specific neuronal subtypes. **(a)** Schematic representation of experimental design that applies to **b-e**. Human brain organoids were SARS-CoV-2-infected at multiplicity of infection 1 for 5 days and subsequently exposed to the indicated senolytic treatments for 5 additional days. Analysis was performed at the end time point of the 10-day experiment. **(b)** SA-β-gal activity was evaluated at 10 days post infection. Bar graphs show the percentage of SA-β-gal-positive cells. Each data point in the bar graph represents a single organoid section analysed. Error bars represent s.d.; at least 5 individual organoids were analysed per variant-infected condition; one-way ANOVA with multiple-comparison post-hoc corrections. Scale bar, 0.3 mm. **(c)** Total RNA from individual organoids uninfected or infected with the SARS-CoV-2 Delta variant was used to quantify the RNA expression levels of the indicated viral genes and normalized to RPLP0 mRNA and compared to infected vehicle controls. Error bars represent s.e.m.; n = 3 independent organoids; one-way ANOVA with multiple-comparison post-hoc corrections; nd: not detected. **(d)** Stacked bars show NanoString GeoMx deconvolved p16-positive ROI cell abundance using constrained log-normal regression from organoids uninfected or infected with the SARS-CoV-2 Delta variant. L4/5/6 IT Car3: Glutamatergic neurons; L5 ET: Cortical layer 5 pyramidal neurons; L6CT L6b: Corticothalamic (CT) pyramidal neurons in layer 6; CGE: GABAergic ganglionic eminence neurons; EC: Endothelial cells; VLMC: vascular and leptomeningeal cells. **(e)** Bar graphs show the percentage of deconvolved p16-positive neuronal populations significantly modulated upon SARS-CoV-2 Delta variant infection and subsequent senolytic interventions. n = 3 independent ROIs per condition tested; *P < 0.05; one-way ANOVA with multiple-comparison post-hoc corrections.

### Senolytic treatments mitigate COVID-19 brain pathology *in vivo*

To investigate the consequences of CNS SARS-CoV-2 infection and ensuing brain virus-induced senescence in a more physiologically complete system, we utilised transgenic mice expressing human ACE2 gene under the control of the keratin 18 promoter (K18-hACE2)^37^ and performed intranasal SARS-CoV-2 infections, where we found brain viral nucleocapsid antigen in cerebral cortex and brainstem regions (Supplementary Fig. 5a). Experimentally, 24 hours post infection we initiated oral administration of the senolytic interventions navitoclax, fisetin and D+Q – drugs known to exert blood-brain barrier permeability^22,38^ – with subsequent treatments every two days (Fig. 6a). As previously reported, SARS-CoV-2-infected K18-hACE2 transgenic mice undergo dramatically shortened lifespans upon infection^37^, with a median survival of 5 days. Strikingly, treatment with D+Q or fisetin significantly improved the survival of K18-hACE2 mice as compared to vehicle-treated controls, with extended median lifespans of 60% (Fig. 6b). Furthermore, while at 10 days post infection all vehicle-treated control mice were already dead, at survival experimental endpoint (12 days post infection) a percentage of senolytic-treated mice – 22% (fisetin), 38% (D+Q) and 13% (navitoclax) – remained alive (Fig. 6b). This significantly improved survival upon senolytic administration of infected mice concurrently delayed the rapid weight loss observed in the infected control group (Supplementary Fig. 5b). Throughout the first week of the *in vivo* experiments, mice were clinically monitored and scored daily for behavioural and physical performance (Fig. 6c). Notably, senolytic interventions resulted in a profound reduction of COVID-related disease features, especially in the D+Q-treated group (Fig. 6c). Given the positive survival and improved clinical performance outcomes brought about by senolytic treatment, we investigated whether the oral administration of senolytics impacted the histological architecture and pro-inflammatory makeup of brains from infected mice. To this end, we first tested the impact of senolytics on brain viral RNA levels. In accordance with our brain organoid data (Fig. 5c), senolytic treatments of infected K18-hACE2 mice showed a significantly lower viral gene expression compared to infected vehicle-treated mice (Fig. 6d), further supporting a putative role for senescent cells in preferentially sustaining SARS-CoV-2 replication. We next tested whether senescent cell clearance directly impacted the transcription of SASP and senescence genes in the brain. mRNA expression analyses from brains of uninfected and infected mice indicated an overall increase in inflammatory SASP and p16 senescence markers in the brains of infected mice (Fig. 6e). Most importantly, all three senolytic interventions consistently normalised brain SASP and senescence gene expression of infected mice to levels comparable to those of uninfected brains (Fig. 6e).

**Figure 6.**
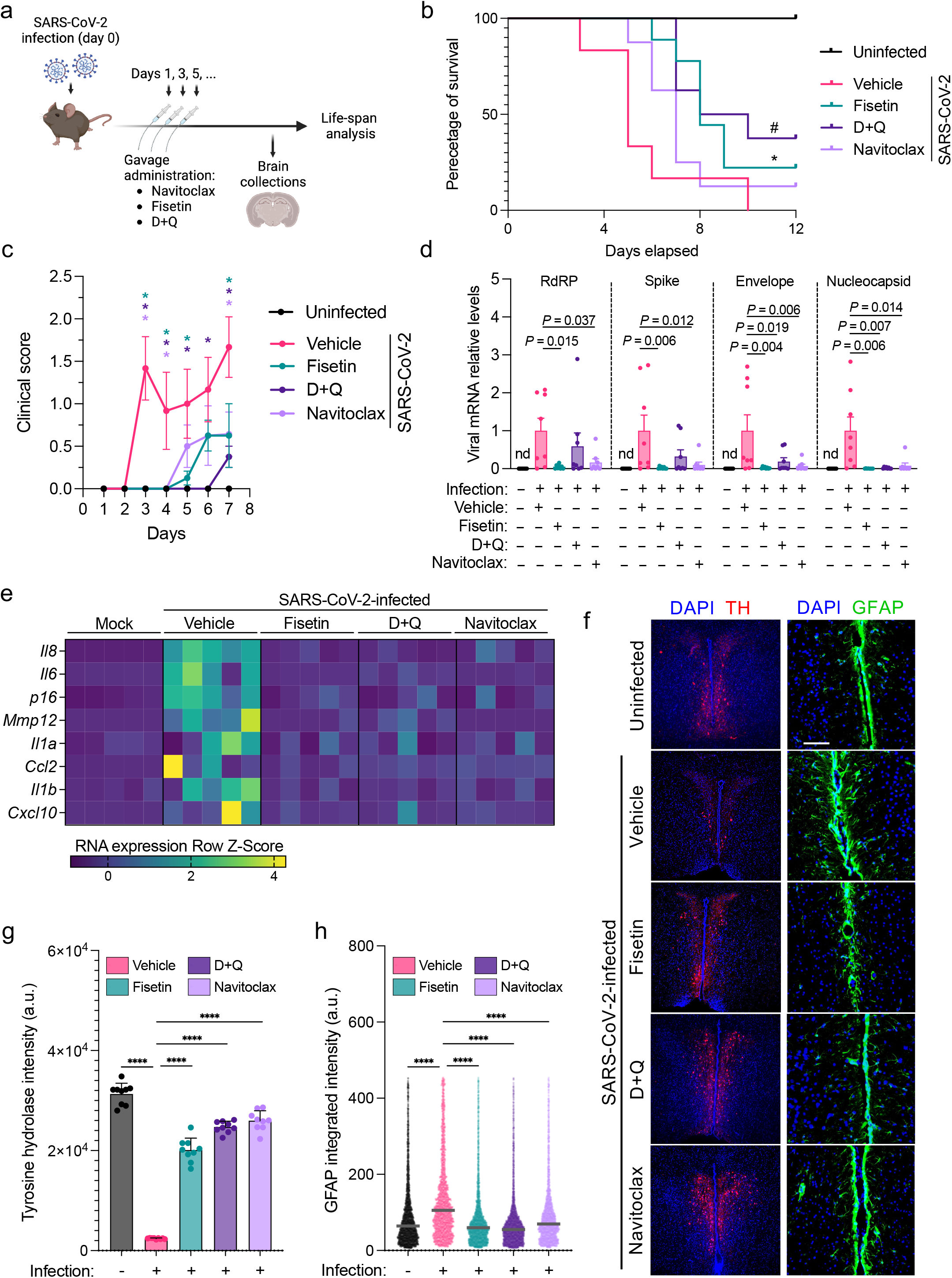
Senolytic treatments mitigate COVID-19 brain pathology *in vivo*. **(a)** Schematic representation of experimental design that applies to **b-h**. K18-hACE2 transgenic mice were exposed to Delta variant infections on day 0 and subsequently treated with the indicated senolytics every other day starting on day 1. Two mouse cohorts were randomly allocated for scheduled euthanasia on day 5 for brain tissue characterisation as well as end time point experiments to monitor clinical score and survival. **(b)** Kaplan–Meier curve of uninfected mice (n = 3), and SARS-CoV-2-infected mice treated with vehicle (n = 6), fisetin (n = 9), D+Q (n = 8), or navitoclax (n = 8). * P = 0.032 for vehicle vs fisetin curve comparison; # P = 0.0087 for vehicle vs D+Q curve comparison; Kaplan–Meier survival analysis. **(c)** Graph shows the average combined behavioural and physical clinical score over time of uninfected mice (n = 3), and SARS-CoV-2-infected mice treated with vehicle (n = 6), fisetin (n = 8), D+Q (n = 8), or navitoclax (n = 8). Error bars represent s.e.m.; color-coded *P<0.05 for comparisons between vehicle and each color-coded senolytic treatment; one-way ANOVA with multiple-comparison post-hoc corrections for every timepoint tested. **(d)** Total RNA of individual brains from mice uninfected or infected with the SARS-CoV-2 Delta variant and treated with various senolytic interventions was used to quantify the RNA expression levels of the indicated viral genes and was normalized to *Rplp0* mRNA and compared to infected vehicle controls. Error bars represent s.e.m.; n = 8 mouse brains per condition; one-way ANOVA with multiple-comparison post-hoc corrections; nd: not detected. **(e)** Total RNA of individual brains from mice uninfected or infected with the SARS-CoV-2 Delta variant and treated with various senolytic interventions was used to quantify the mRNA expression levels of the indicated senescence and SASP genes and was normalized to *Rplp0* mRNA. Each column in the heatmap represents an individual mouse brain analysed. **(f)** Representative immunofluorescent images of brainstem regions of coronal brain sections from uninfected or infected mice with the SARS-CoV-2 Delta variant and treated with the indicated senolytics. Samples were immunolabelled with antibodies against TH (red; scale bar, 100 μm) and GFAP (green; scale bar, 50 μm). **(g)** Quantification of the TH data presented in **f.** Bar graph shows the intensity of tyrosine hydrolase (TH) staining within the brainstem. Each data point in the bar graph represents a single brain section analysed. Error bars represent s.d.; ****P<0.0001; 3 brains per condition were analysed; one-way ANOVA with multiple-comparison post-hoc corrections. **(h)** Quantification of the GFAP data presented in **f.** Dot plot shows the intensity of GFAP per cell within the brainstem. Each data point in the dot blot represents a single cell analysed. ****P<0.0001; 3 brains per condition were analysed; one-way ANOVA with multiple-comparison post-hoc corrections.

Neuroinvasive viral infections can result in loss of dopaminergic neurons and ensuing PASC such as parkinsonism^39^. Given the long-term neurological impact of COVID-19 including coordination and consciousness disorders^40^, we therefore tested the impact of SARS-CoV-2 infection on altering dopaminergic neuron survival within the brainstem, an important region of the brain known to regulate these behaviours. Strikingly, Delta variant infection induced a dramatic loss of dopaminergic neuronsin the brainstem, as measured by tyrosine hydroxylase immunolabelling (Fig. 6f, g), and this was accompanied by increased astrogliosis (Fig. 6f, h), a neurotoxic process common to multiple neurological disorders^41^. Importantly, recurrent senolytic treatments initiated 24 hours after SARS-CoV-2 exposure partly prevented dopaminergic neuron loss and abrogated the onset of reactive astrogliosis (Fig. 6f-h).

## Discussion

Brain aging and related cognitive deficiency have been attributed to diverse molecular processes including chronic inflammation and cellular senescence^42^. This has been studied both in normal murine aging^24^, as well as in different age-related mouse models of neurodegeneration such as Parkinson’s disease^43^, tauopathies^23,44^, amyloid-beta neuropathology^22^, and neuropsychiatric disorders^45^. However, whether the endogenous age-related onset of cellular senescence impacts brain aging in human tissue systems has not been investigated. Neither have the putative consequences of neurotropic viral infections in accelerating the onset of cellular senescence in the brain been examined.

Our findings herein show that: (1) senescent cells accumulate in physiologically aged brain organoids of human origin and that long-term (4 weeks), intermittent, senolytic treatment reduces inflammation and cellular senescence; (2) interventions unique to D+Q treatments induce antiaging and pro-longevity gene expression changes in human BOs; (3) brains from COVID-19 patients undergo accelerated cellular senescence accumulation compared to age-matched controls; (4) SARS-CoV-2 and neurotropic viruses including Zika and JEV can infect human BOs to directly induce cellular senescence; (5) Delta (B.1.617.2) variant induces the strongest SARS-CoV-2-dependent induction of cellular senescence, where spatial transcriptomic sequencing of p16-positive cells identifies a Delta-specific SASP signature; (6) short-term (5 days) senolytic treatments of SARS-CoV-2-infected organoids reduce viral gene expression and prevent the onset of senescent neurons of corticothalamic and GABAergic nature; and (7) senolytic treatment following SARS-CoV-2 intranasal infection of K18-hACE2 mice ameliorates COVID-19 neuropathology, including improvements in clinical score and survival, alleviation of reactive astrogliosis, increased survival of dopaminergic neurons, and reduced viral, SASP and senescence gene expression in the brain of infected mice.

To evaluate the relationship between senescent cell accumulation and brain aging, we designed studies to eliminate senescent cells through pharmacologic approaches (D+Q, navitoclax and ABT-737) and hypothesized that senolytic interventions may have beneficial consequences in targeting brain aging. We found that physiologically aged human BOs accumulate senescent cells and that senolytic treatment can be used as a proof-of-concept strategy to revert Lamin B1 levels, and alleviate differential SASP expression and senescent cell burden in human brain BOs systems. In addition to senolytic activity, transcriptomic aging clocks identified D+Q as an intervention that achieved tissue rejuvenation, as 8-month-old human brain organoids displayed comparable aging clocks to D+Q-treated 9-month-old counterparts. Given that senescent cell clearance results in reversal of the aging process, these findings support an important role for senescent cells in driving human brain aging.

Further to normal brain aging, we tested the possibility of virus-induced senescence upon BOs neurotropic infections. We found that flavivirus JEV, ROCV and ZIKV infections, and multiple SARS-CoV-2 variant infections lead to a significant increase in BO cellular senescence. Importantly, upon senolytic delivery BOs display a dramatic loss of SARS-CoV-2 viral RNA expression, suggestive of a role for senescent cells in preferentially facilitating viral entry and retention, consistent with data showing increased ACE2 expression in human senescent cells^46^.

Furthermore, SARS-CoV-2 induces metabolic changes in infected and neighbouring neurons^8^, a paracrine phenomenon reminiscent of the bystander effect characteristic of senescent cells^47^. Here, spatial transcriptomic sequencing cell deconvolution of p16 protein-expressing cell clusters identified two neuronal populations – corticothalamic and GABAergic – that become senescent and broadly develop a *de novo* SASP signature upon Delta (B.1.617.2) infection. It will therefore be of interest to determine whether neuronal virus-induced senescence contributes to neuroinflammation and the long-term neurological impact of COVID-19.

In the brains of SARS-CoV-2-infected K18-hACE2 mice, we found that senolytic treatment alleviates p16 and the levels of proinflammatory cytokines which may be due, in part, to removal of virus-induced senescence and ensuing SASP expression. However, secondary anti-inflammatory and/or anti-viral effects of D+Q, fisetin or navitoclax – for instance by direct inhibition of the observed astrogliosis – are also possible. Upon systematic monitoring of clinical performance in SARS-CoV-2-infected mice, we found that intermittent senolytic treatment significantly improved animal behaviour and survival. This beneficial clinical effect of senolytics was associated with reduced inflammation and increased survival of dopaminergic neurons. Indeed, inflammatory cytokines as part of the SASP can impair brain plasticity^48^, suggesting that the beneficial effects of senolytic treatment on COVID-19 neurological clinical picture may result from suppression of senescence-dependent inflammation and improved neuronal survival. Whether the *in vivo* effects of senolytics on COVID-19 neuropathology exclusively results from clearance of cellular senescence or also involves actions on dopaminergic neurons and other brain regions remains to be determined. Nevertheless, in this study we have provided important evidence that paves the way for future preclinical and clinical studies that will test the hypothesis that senolytic therapies can suppress long-COVID neuropathology and other long-term disorders caused by acute neurotropic viral infections.

## Methods

### Ethics and biological safety

The use of animals was approved by the University of Queensland Animal Ethics Committee under project number 2021/AE001119. Mice were housed within the BSL-3 facility using IsoCage N-Biocontainment System (Tecniplast, USA), where each cage was supplied with a HEPA filter, preventing viral contamination between cages. This IsoCage system also provides individual ventilation to the cages, maintaining the humidity under 65-70% and temperature between 20–23 °C. Mice were kept under a 12-h light/dark cycle with food and water provided ad libitum.

Pathogenic SARS-CoV-2 variants and encephalitic flaviviruses were handled under a certified biosafety level-3 (BSL-3) conditions in the School of Chemistry and Molecular Biosciences (SCMB), Australian Institute for Bioengineering and Nanotechnology (AIBN) and Institute for Molecular Bioscience (IMB) at The University of Queensland, Australia. All approved researchers have used disposal Tychem 2000 coveralls (Dupont, Wilmington, NC, USA; #TC198T YL) at all times and used powered air-purifying respirators (PAPR; SR500 Fan Unit) or Versaflo-powered air-purifying respirators (3M, Saint Paul, MN, USA; #902-03-99) as respiratory protection. All pathogenic materials were handled in a class II biosafety cabinet within the BSL-3 facility. For downstream analysis, all samples containing infectious viruses were appropriately inactivated in accordance with the BSL-3 manual. Liquid and solid waste were steam-sterilised by autoclave. This study was approved by the Institutional Biosafety Committee from The University of Queensland (UQ) under the following approvals IBC/485B/SCMB/2021 and IBC/447B/SCMB/2021. The analysis of human brain sections was performed with the approval of the Ethic Committee of the University of Freiburg: 10008/09. The study was performed in agreement with the principles expressed in the Declaration of Helsinki (2013).

### Generation and culture of PSC-derived human brain organoids

Organoid generation was carried out as previously described^49^. Human H9 (WA09) pluripotent stem cells (hPSCs) were obtained from WiCell with verified normal karyotype and contamination-free; and were routinely tested and confirmed negative for mycoplasma (MycoAlert, Lonza). hPSCs were maintained in mTeSR media (STEMCELL Technologies, cat. #85850) on matrigel-coated plates (Corning, No. 354234). On day 0 of organoid differentiation, PSCs were dissociated with Accutase (Life Technologies, cat. #00-4555-56) and seeded at a density of 15,000 cells per well on a 96-well low-attachment U-bottom plate (Sigma, cat. #CLS7007) in mTeSR plus 10 μM ROCK inhibitor (VWR, cat. #688000-5). The 96 well-plate was then spun at 330 g for 5 minutes to aggregate the cells and make spheroids. The spheroids were fed every day for 5 days in media containing Dulbecco’s modified eagle medium (DMEM)/F12 (Invitrogen, cat. #11330-032), Knock-out serum (Invitrogen, cat. #11320-033), 1:100 Glutamax, 1:200 MEM-NEAA supplemented with dual SMAD inhibitors: 2 μM Dorsomorphin (StemMACS, cat. #130-104-466) and 2 μM A-83-01 (Lonza, cat. #9094360). On day 6, half of the medium was changed to induction medium containing DMEM/F12, 1:200 MEM-NEAA, 1:100 Glutamax, 1:100 N2 supplement (Invitrogen, cat. #17502048), 1 μg ml-1 heparin (Sigma, cat. # H3149) supplemented with 1 μM CHIR 99021 (Lonza, cat. #2520691) and 1 μM SB-431542 (Sigma, cat. # S4317). From day 7, complete media change was done with induction media followed by everyday media changes in induction media for the next 4 days. On day 11 of the protocol, spheroids were transferred to 10 μl-droplets of Matrigel on a sheet of Parafilm with small 2 mm dimples. These droplets were allowed to gel at 37°C for 25 minutes and were subsequently removed from the Parafilm and transferred to and maintained in low-attachment 24-well plates (Sigma, cat. #CLS3473) containing induction medium for the following 5 days. From day 16, the medium was then changed to organoid medium containing a 1:1 mixture of Neurobasal medium (Invitrogen, cat. #21103049) and DMEM/F12 medium supplemented with 1:200 MEM-NEAA, 1:100 Glutamax, 1:100 N2 supplement, 1:50 B27 supplement (Invitrogen, cat. #12587010), 1% penicillin-streptomycin (Sigma, cat. #P0781), 50 μM 2-mercaptoethanol and 0.25% insulin solution (Sigma, cat. #I9278). Media was changed every other day with organoid medium. Organoids were maintained in organoid media until the end of experiments, as indicated.

### Human tissue preparation

frontal cortex tissue from patients that had tested positive for SARS-CoV-2 and died from severe COVID-19 was obtained at the University Medical Center Freiburg, Germany. The tissue was formalin-fixed and embedded into paraffin (FFPE) using a Tissue Processing Center (Leica ASP300, Leica). Sections (3 μm thick) were cut and mounted onto Superfrost objective slides (Langenbrinck).

### Cell lines

RNA Vero E6 cells (African green monkey kidney cell clones) and TMPRSS2-expressing Vero E6 cell lines were maintained in Dulbecco’s Modified Eagle Medium (DMEM, Gibco, USA) at 37 °C with 5 % CO2. Additionally, as previously described, the TMPRSS2-expressing Vero E6 cell line was supplemented with 30 μg/mL of puromycin^50^. C6/36 cells, derived from the salivary gland of the mosquito *A. albopictus* were grown at 28 °C in Royal Park Memorial Institute (RPMI) medium (Gibco, USA). All cell lines media were supplemented with 10% heat-inactivated foetal calf serum (FCS) (Bovogen, USA), penicillin (100 U/mL) and streptomycin (100 μg/mL) (P/S). C6/36 media was also supplemented with 1% GlutaMAX (200 mM; Gibco, USA) and 20 mM of HEPES (Gibco, USA). All cell lines used in this study were tested mycoplasma free by first culturing the cells for 3-5 days in antibiotic-free media and then subjected to a mycoplasma tested using MycoAlert™ PLUS Mycoplasma Detection Kit (Lonza, UK).

### Viral isolates

Seven SARS-CoV-2 variants were used in this study. *i*) Ancestral or Wuhan strain: an early Australian isolate hCoV-19/Australia/QLD02/2020 (QLD02) sampled on 30/01/2020 (GISAID Accession ID; EPI_ISL_407896); *ii*) Alpha (B.1.1.7) named as hCoV-19/Australia/QLD1517/2021 and collected on 06/01/2021 (GISAID accession ID EPI_ISL_944644); *iii*) Beta (B.1.351), hCoV19/Australia/QLD1520/2020, collected on 29/12/2020 (GISAID accession ID EPI_ISL_968081); *iv*) Delta (B.1.617), hCoV-19/Australia/QLD1893C/2021 collected on 05/04/2021 (GISAID accession ID EPI_ISL_2433928); *v*) Gamma (P.1), hCoV-19/Australia/NSW4318/2021 sampled on 01-03-2021 (GISAID accession ID EPI_ISL_1121976); *vi*) Lambda (C.37), hCoV-19/Australia/NSW4431/2021 collected on 03-04-2021 (GISAID accession ID EPI_ISL_1494722); and *vii*) Omicron (BA.1), hCoV-19/Australia/NSW-RPAH-1933/2021 collected on 27-11-2021 (GISAID accession ID EPI_ISL_6814922). All viral isolates obtained were passaged twice except for Gamma and Lambda variants, which were passed thrice. Viral stocks were generated on TMPRSS2-expressing Vero E6 cells to ensure no spike furin cleavage site loss. To authenticate SARS-CoV-2 isolates used in the study viral RNA was extracted from stocks using TRIzol LS reagent (Thermo Fisher Scientific, USA) and cDNA was prepared with Protoscript II first-strand cDNA synthesis kit as per manufacturer’s protocol (New England Biolabs, USA). The full-length Spike glycoprotein was subsequently amplified with Prime Star GXL DNA polymerase (Takara Bio) and the following primers CoV-SF GATAAAGGAGTTGCACCAGGTACAGCTGTTTTAAG CoV-SR GTCGTCGTCGGTTCATCATAAATTGGTTCC and conditions as per previously described^50^. For encephalitic flaviviruses, virulent strains of Zika virus (ZIKV, Natal [GenBank: KU527068.1]), Japanese encephalitis virus (JEV, Nakayama strain [GenBank: EF571853.1]) and Rocio virus (ROCV, [GenBank: AY632542.4]) were propagated on C6/36 to generate a viral stock for all the experiments. Viral titres were determined by an immuno-plaque assay^51^.

### RNA isolation

RNA from brain organoids and mouse tissue was extracted with RNeasy Mini Kit (Qiagen) for mRNA detection, according to the manufacturer’s instructions. Mouse tissue was homogenised with a TissueLyser II (Qiagen) at 30 Hz for 60 seconds. RNA integrity of brain organoids and mouse tissue was evaluated by analysis on the 2100 Bioanalyzer RNA 6000 Pico Chip kit (Agilent) using the RNA Integrity Number (RIN). RNA samples with a RIN > 7 were considered of high enough quality for real-time quantitative PCR, and transcriptomic library construction and RNA sequencing according to the manufacturer’s instructions.

### Real-time quantitative PCR

1 μg of total RNA was reverse transcribed using iScript cDNA Synthesis Kit (Bio-Rad). A volume corresponding to 5 ng of initial RNA was employed for each real-time PCR reaction using PowerUp SYBR Green Master Mix (Applied Biosystems) on a CFX Opus Real-Time PCR detection system. Ribosomal protein P0 (RPLP0) were used as control transcripts for normalization. Primers sequences (5’-3’ orientation) are listed in Supplementary Table 1.

### Viral infection of organoids

Brain organoids in low-adhesion plates were infected overnight (14 hours) with the indicated flaviviruses and SARS-CoV-2 variants at multiplicity of infection (MOI) 0.1 and 1, respectively. Then, brain organoids were thrice washed with LPS-free PBS and added maintenance media and kept for 5 days post-infection.

### Senolytic treatments *in vitro*

For infection experiments, 5 days after viral exposure brain organoids were treated with a single dose of navitoclax (2.5 μM), ABT-737 (10 μM) or D+Q (D: 10 μM; Q: 25 μM) and monitored for 5 days following treatment. As for senolytic interventions on physiologically aged 8-month-old organoids, brain organoids were treated with a weekly dose of navitoclax (2.5 μM), ABT-737 (10 μM) or D+Q (D: 10 μM; Q: 25 μM) for 4 weeks and subsequently collected for downstream analysis.

### SARS-CoV-2-driven COVID-19 animal experiments

*In vivo* experiments were performed using 6-week-old K18-hACE2 transgenic female mice obtained from the Animal Resources Centre (Australia). For animal infections, SARS-CoV-2 was delivered intranasally — 20 μl of the Delta variant at 5 x 10^3^ FFU per mouse — on anesthetized mice (100 mg kg^-1^ ketamine and 10 mg kg^-1^ xylazine). Control animals were mock-infected with the same volume of RPMI additive-free medium. One day after infection, K18-hACE2 mice were randomly distributed into three treatment groups (n = 16 each) and one solvent-only control group (n = 16). From 1 day after infection, randomly chosen animals were treated via oral gavage routes with navitoclax (100 mg kg^-1^), D+Q (D: 5 mg kg^-1^; Q: 50 mg kg^-1^) or fisetin (100 mg kg^-1^) dissolved in 5% DMSO and 95% corn oil every other day. For tissue characterization (n = 8 for each infected group), on day 6 after infection animals were euthanised and brain specimens were collected for RNA expression analysis and histopathological assessment. For clinical and survival evaluation, mice were monitored daily for up to 12 days post infection. Clinical scoring included: no detectable disease (0); hindlimb weakness, away from littermates, ruffled fur (0.5-1); partial hindlimb paralysis, limping, hunched, reluctant to move (1.5-2); and complete paralysis of hindlimb, severely restricted mobility, severe distress, or death (2.5-3).

### Organoid sectioning and histology

Brain organoids were fixed in 4% paraformaldehyde (PFA) for 1 hour at RT and washed with phosphate-buffered saline (PBS) three times for 10 minutes each at RT before allowing to sink in 30% sucrose at 4°C overnight and then embedded in OCT (Agar Scientific, cat. #AGR1180) and cryosectioned at 14 μm with a Thermo Scientific NX70 Cryostat. Tissue sections were used for immunofluorescence and for the SA-β-Gal assay. For immunofluorescence, sections were blocked and permeabilized in 0.1% Triton X-100 and 3% Bovine Serum Albumin (BSA) in PBS. Sections were incubated with primary antibodies overnight at 4°C, washed and incubated with secondary antibodies for 40 minutes at RT. 0.5 μg ml-1 DAPI (Sigma, cat. #D9564) was added to secondary antibody to mark nuclei. Secondary antibodies labelled with Alexafluor 488, 568, or 647 (Invitrogen) were used for detection. SA-β-gal activity at pH 6.0 as a senescence marker in fresh or cryopreserved human samples was assessed as previously described^52^.

### Nanostring spatial transcriptomics

OCT-embedded organoids were freshly sectioned and prepared according to the GeoMX Human Whole Transcriptome Atlas Assay slide preparation for RNA profiling (NanoString). Fastq files were uploaded to GeoMX DSP system where raw and Q3 normalized counts of all targets were aligned with ROIs. Cell abundances were estimated using the SpatialDecon R library, which performs mixture deconvolution using constrained log-normal regression. The 0.75 quantile-scaled data was used as input. DESeq2 R package^53^ was used to identify differently expressed genes in the ROI cell subsets. DESeq2 was performed between the pairwise comparisons of interest and genes were corrected using the Benjamini & Hochberg correction and only genes that had a corrected P-value of < 0.05 were retained.

### Whole organoid RNA sequencing

Before mRNA sequencing, ribosomal RNA from bulk organoid RNA was depleted with the Ribo-Zero rRNA Removal Kit (Illumina). Transcripts were sequenced at Novogene Ltd (Hong Kong) using TruSeq stranded total RNA library preparation and Illumina NovaSeq 150bp paired-end lane. FastQC was used to check quality on the raw sequences before analysis to confirm data integrity. Trimmed reads were mapped to the human genome assembly hg38 using Hisat2 v2.0.5. To ensure high quality of the count table, the raw count table generated by featureCounts v1.5.0-p3 was filtered for subsequent analysis. Differential gene expression analysis was performed using Bioconductor DESeq2 R packages. The resulting P-values were adjusted using the Benjamini and Hochberg’s approach for controlling the false discovery rate. Genes with an adjusted P-value <0.05 found by DESeq2 were assigned as differentially expressed.

### Association with gene expression signatures of aging and longevity

To assess the effect of senolytics on transcriptomic age of BO samples, we applied brain multi-species (mouse, rat, human) transcriptomic clock based on signatures of aging identified as explained in^54^. The missing values were omitted with the precalculated average values from the clock. Association of gene expression log-fold changes induced by senolytics in aged BO with previously established transcriptomic signatures of aging and established lifespan-extending interventions was examined as described in^54^. Utilized signatures of aging included multi-species brain signature as well as multi-tissue aging signatures of mouse, rat and human. Signatures of lifespan-extending interventions included genes differentially expressed in mouse tissues in response to individual interventions, including caloric restriction (CR), rapamycin (Rapamycin), and mutations associated with growth hormone deficiency (GH deficiency), along with common patterns of lifespan-extending interventions (Common) and ECs associated with the intervention effect on mouse maximum (Max lifespan) and median lifespan (Median lifespan).

For the identification of enriched functions affected by senolytics in aged BO we performed functional GSEA^55^ on a pre-ranked list of genes based on log10(p-value) corrected by the sign of regulation, calculated as:

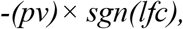

where pv and lfc are p-value and logFC of a certain gene, respectively, obtained from edgeR output, and sgn is the signum function (equal to 1, −1 and 0 if value is positive, negative or equal to 0, respectively). HALLMARK ontology from the Molecular Signature Database was used as gene sets for GSEA. The GSEA algorithm was performed separately for each senolytic via the fgsea package in R with 5,000 permutations. A q-value cutoff of 0.1 was used to select statistically significant functions.

Similar analysis was performed for gene expression signatures of aging and lifespan-extending interventions. Pairwise Spearman correlation was calculated for individual signatures of senolytics, aging and lifespan-extending interventions based on estimated NES (Fig. 2g). A heatmap colored by NES was built for manually chosen statistically significant functions (adjusted p-value < 0.1) (Supplementary Fig. 1a).

### Imaging and analysis

Immunofluorescence images were acquired using a Zeiss LSM 900 Fast Airyscan 2 super-resolution microscope or a Zeiss AxioScan Z1 Fluorescent Imager. For organoid staining, the number of positive cells per organoid for senescence, cell type and viral markers tested was analysed by the imaging software CellProfiler, using the same pipeline for each sample in the same experiment. Custom Matlab scripts were developed to streamline high throughput imaging data.

### Antibodies

anti-p16 (Cell Signalling, 80772, 1:400); anti-NeuN (Millipore, ABN78, 1:1000); anti-GFAP (Agilent, Z0334, 1:2000); anti-Sox2 (Cell Signalling, 23064, 1:1000); anti-SARS-CoV-2 Nucleocapsid C2^56^; anti-SARS-CoV-2 spike protein^57^; anti-γH2AX (Millipore, 05-636, 1:1000); anti-Tyrosine Hydroxylase (Invitrogen, PA5-85167, 1:1000); anti-lamin B1 (Abcam, ab16048, 1:5000); anti-Chicken IgG (Jackson ImmunoResearch, 703-545-155, 1:500); anti-rabbit IgG (Invitrogen, A10042, 1:400); anti-rabbit IgG (Invitrogen, A21245, 1:400); anti-mouse IgG (Invitrogen, A11029, 1:400); anti-mouse IgG (Invitrogen, A21235, 1:400); anti-human IgG (Invitrogen, A21445, 1:400).

### Statistical analysis

Results are shown as mean ± standard error of the mean (s.e.m.) or standard deviation (s.d.) as indicated. No statistical methods were used to predetermine sample size. P value was calculated by the indicated statistical tests, using R or Prism software. In figure legends, n indicates the number of independent experiments or biological replicates.

## Data availability

RNA-seq raw data are being deposited in the European Nucleotide Archive. RNA-seq files from Mavrikaki et al. are available through the Gene Expression Omnibus accession number GSE188847.

## Acknowledgements

We thank Novogene for performing bulk RNA sequencing experiments and bioinformatic analysis; Aaron McClelland from NanoString (Seattle, USA) for technical and computational assistance on GeoMx spatial transcriptomic sequencing; the scientists and pathologists of Queensland and New South Wales Department of Health, and Kirby Institute for providing the SARS-CoV-2 variants; Maya Patrick and Barb Arnts (UQBR animal staff) at the AIBN and Crystal McGirr (BSL-3 facility manager at the IMB) for technical assistance; Robert Sullivan from the Queensland Brain Institute for technical advice; Jasmyn Cridland (Regulatory Compliance Officer, Faculty of Science at UQ) and Amanda Jones (UQ Biosafety) for advice on Biosafety approvals and BSL-3 manual and safety procedures; Shaun Walters, David Knight and Erica Mu from the School of Biomedical Sciences Imaging and Histology facilities (The University of Queensland) for technical support; and EW, JM and DW laboratory members for discussions. MS was supported by the Berta-Ottenstein-Programme for Clinician Scientists, Faculty of Medicine, University of Freiburg, and the IMM-PACT-Programme for Clinician Scientists, Department of Medicine II, Medical Center – University of Freiburg and Faculty of Medicine, University of Freiburg, funded by the Deutsche Forschungsgemeinschaft (DFG, German Research Foundation) – 413517907. TW was supported by the NHMRC (2009957). EW was supported by the NHMRC and an ARC Discovery Project (DP210103401). JA was supported by a University of Queensland Early Career Researcher Grant (application UQECR2058457), a National Health and Medical Research Council (NHMRC) Ideas Grant (2001408), a Brisbane Children’s Hospital Foundation grant (Project-50308) and a Jérôme Lejeune Postdoctoral Fellowship.

## Contributions

JA and HC generated human brain organoids. JA, HC, AT, ATF, MD, MS, AA, GP, EA, NM, BL, AI, DP, IJ, AB, MF, RP, JS, CG, TW, JM and EW contributed to acquisition, analysis, or interpretation of data. AAA, EA, NM and BL participated in the infections and treatments of mice and monitored their clinical performance. JA, ATF and AT analysed transcriptomic data. JA, AA, AF, EA, JM and EW contributed to experimental design. JA planned and supervised the project and wrote the paper. All authors edited and approved the final version of this article.

## Supplementary Figure legends

**Supplementary Figure 1.**
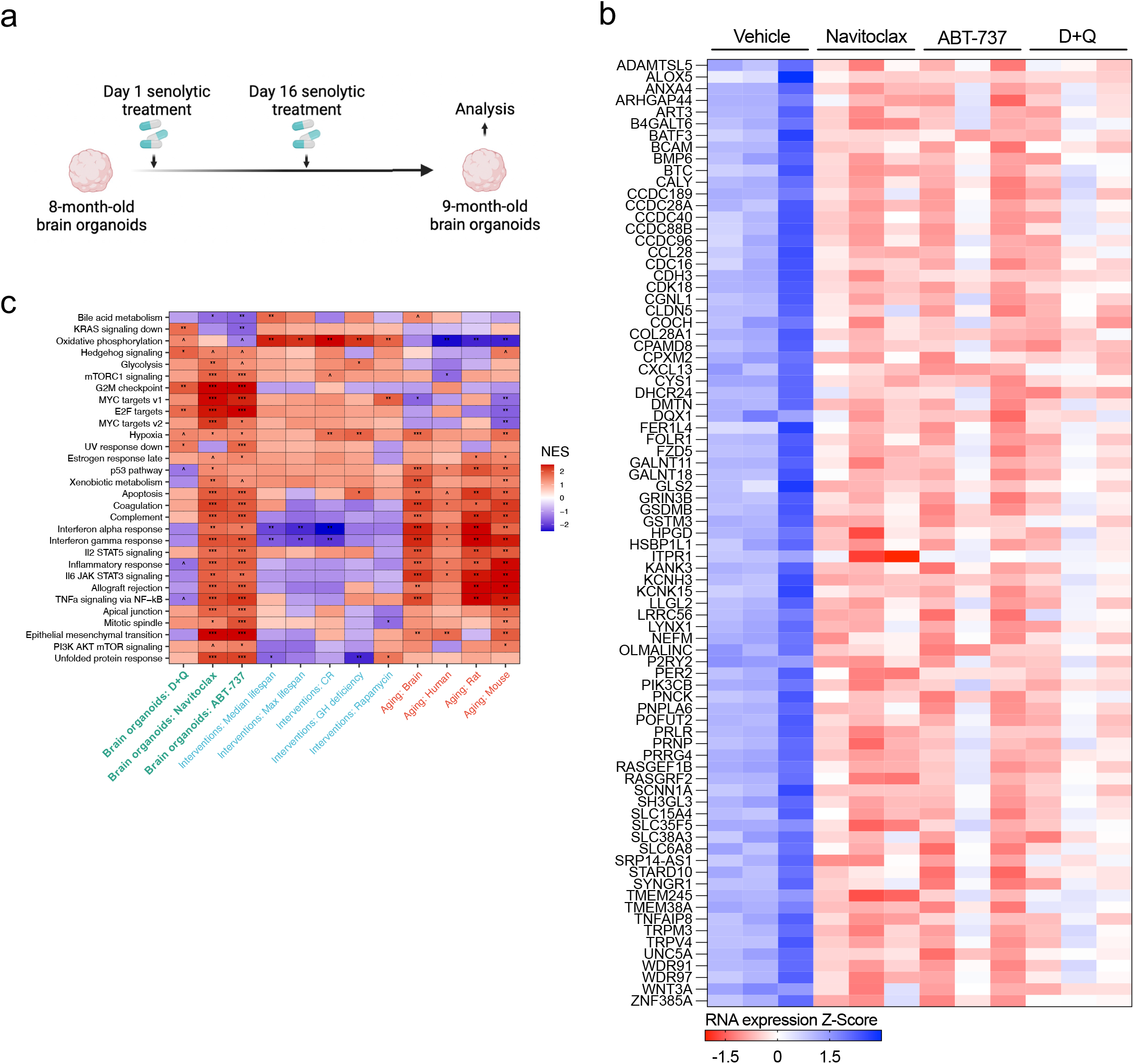
**(a)** Schematic representation of experimental design that applies to Fig. 1,2 and to Supplementary Fig. 1b-c. 8-month-old human brain organoids were exposed to two doses of the senolytic treatments navitoclax (2.5 μM), ABT-737 (10 μM) or D+Q (D: 10 μM; Q: 25 μM): the first one on day 1 and the second dose on day 16. Analysis was performed at the end time point of the 1-month experiment as well as at initial timepoint of 8 months organoid culture. **(b)** Heat map shows senescence-associated RNA transcriptomic expression of downregulated genes shared across all three senolytic interventions. **(c)** Functional enrichment analyses of gene expression signatures and multiple senolytic treatment of brain organoids. Heat map cells are coloured based on the normalized enrichment score (NES).

**Supplementary Figure 2.**
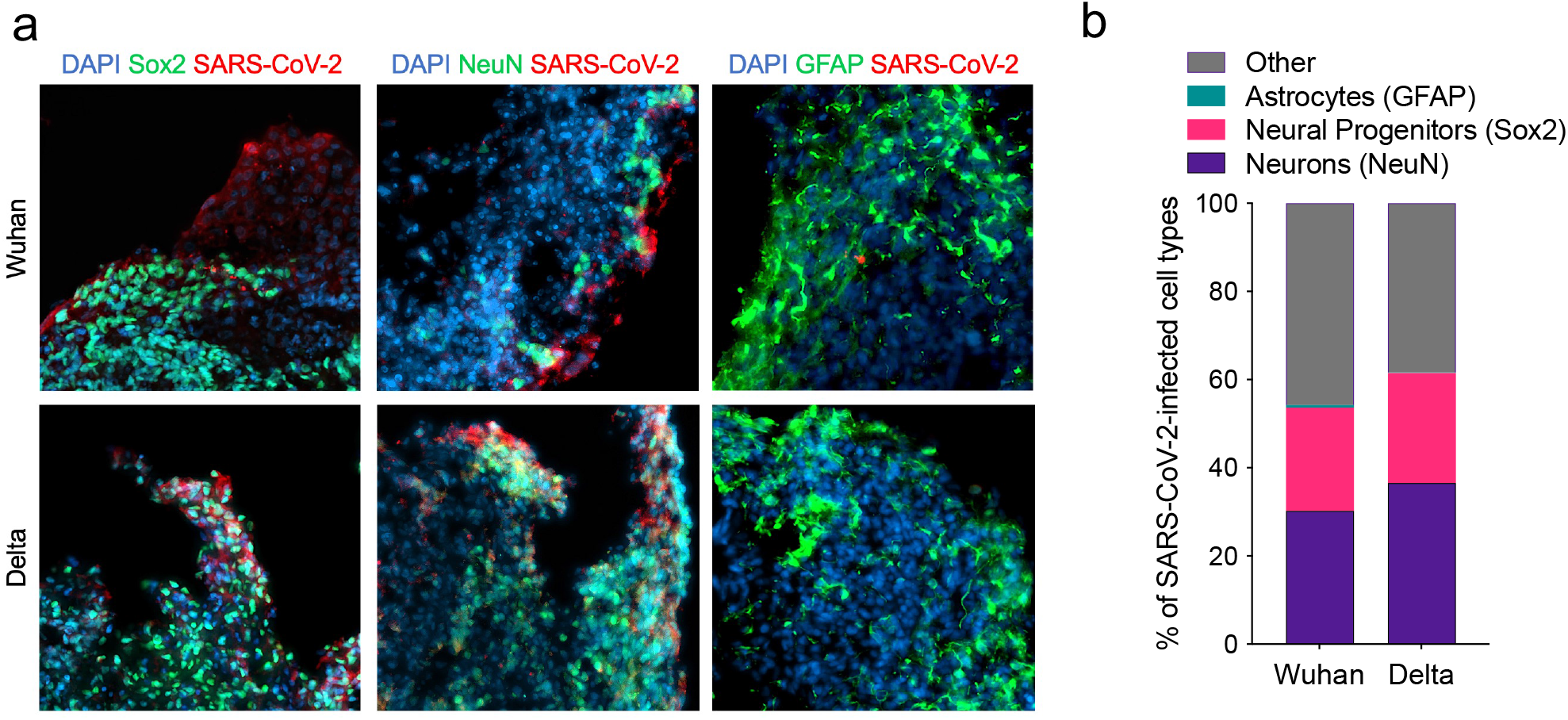
**(a)** Representative images of neural progenitors (Sox2), neurons (NeuN), or astrocytes (GFAP) co-stained with SARS-CoV-2 nucleocapsid protein. Human brain organoids were 3 month-old at time of infection with the indicated SARS-CoV-2 variants at multiplicity of infection 1. **(e)** Stacked bar graphs shoe quantifications from **d**.

**Supplementary Figure 3.**
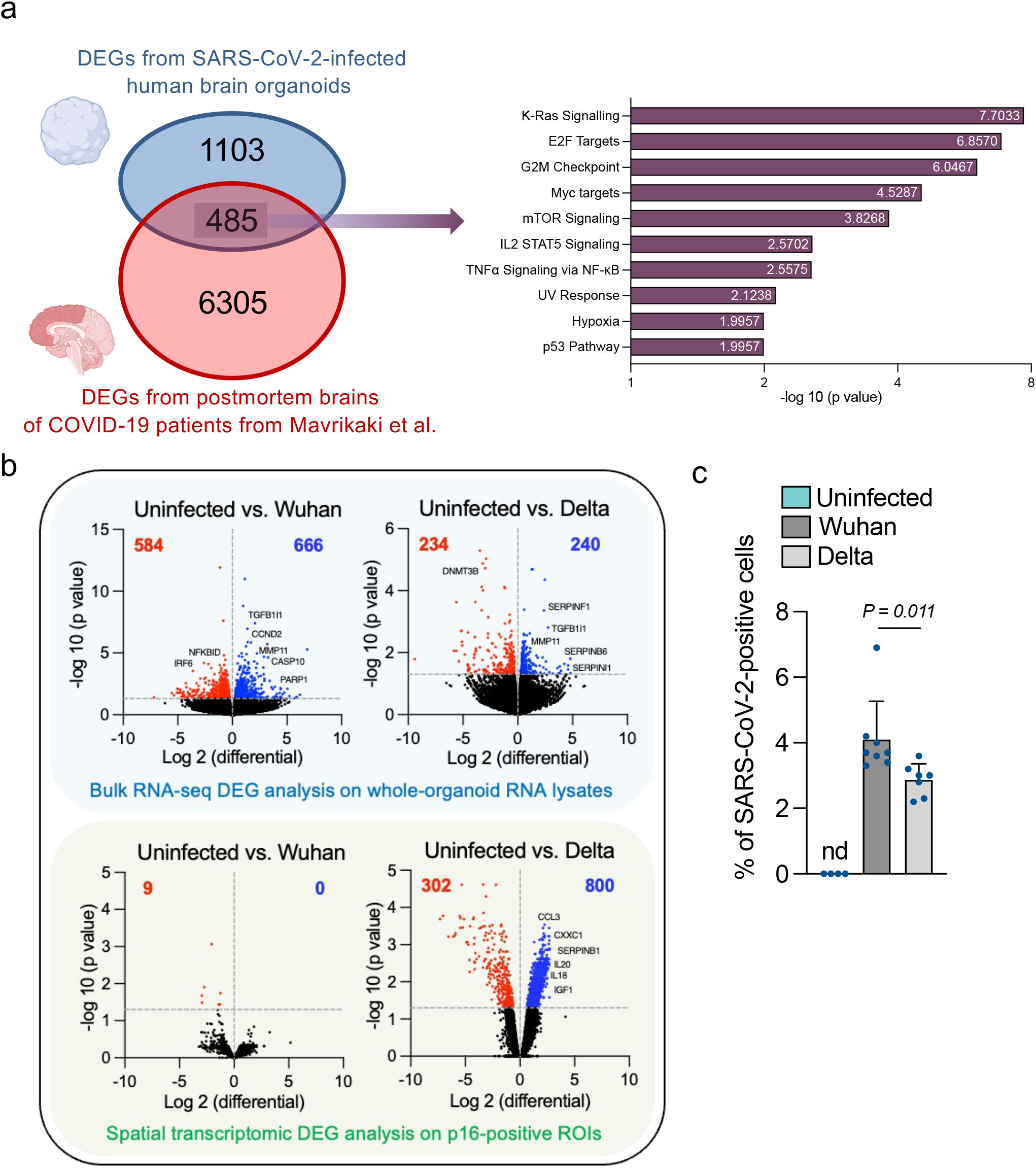
**(a)** Venn diagram on the left shows 485 differentially expressed genes shared across SARS-CoV-2-infected organoids and postmortem brains of COVID-19 patients defined with a significance adjusted P value <0.05. On the right panel, bar graph indicates the pathways enriched within this 485-gene cohort. Gene Set Enrichment Analysis was carried out using aging hallmark gene sets from the Molecular Signature Database. The statistically significant signatures were selected (FDR < 0.25). **(b)** Volcano plots show uninfected versus either Wuhan or Delta-infected brain organoid differential expression of upregulated (blue) and downregulated (red) genes. DEG analysis was performed from whole-organoid RNA-seq data and p16-positive senescent-cell regions of interest (ROIs) from NanoString spatial transcriptomic sequencing. **(c)** Bar graph shows quantifications of nucleocapsid-positive cells from brain organoids infected with the indicated SARS-CoV-2 variants and analysed at 5 days post infection. Each data point in the bar graph represents a single organoid section analysed. Error bars represent s.d.; at least 7 individual organoids were analysed per variant-infected condition; one-way ANOVA with multiple-comparison post-hoc corrections.

**Supplementary Figure 4.**
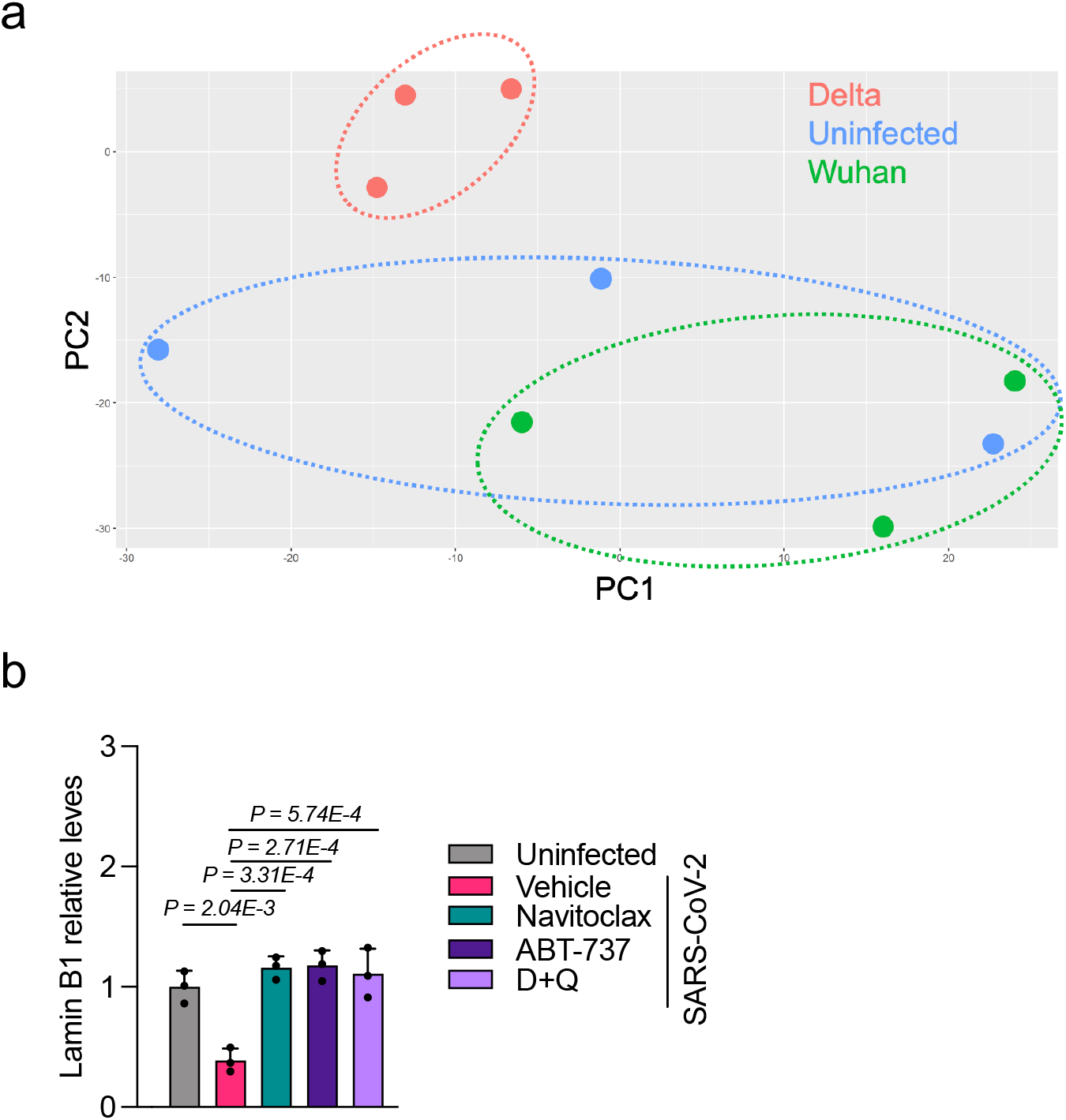
**(a)** Principal component analysis from NanoString spatial transcriptomic sequencing of p16-positive cells in the subspace defined by these differential genes showing clustering of uninfected and Wuhan-infected human brain organoids away from the Delta-infected counterparts. **(b)** Total RNA from individual organoids uninfected or infected with the SARS-CoV-2 Delta variant was used to quantify Lamin B1 mRNA expression levels and normalized to RPLP0 mRNA and compared to infected vehicle controls. Error bars represent s.d.; n = 3 independent organoids; one-way ANOVA with multiple-comparison post-hoc corrections.

**Supplementary Figure 5.**
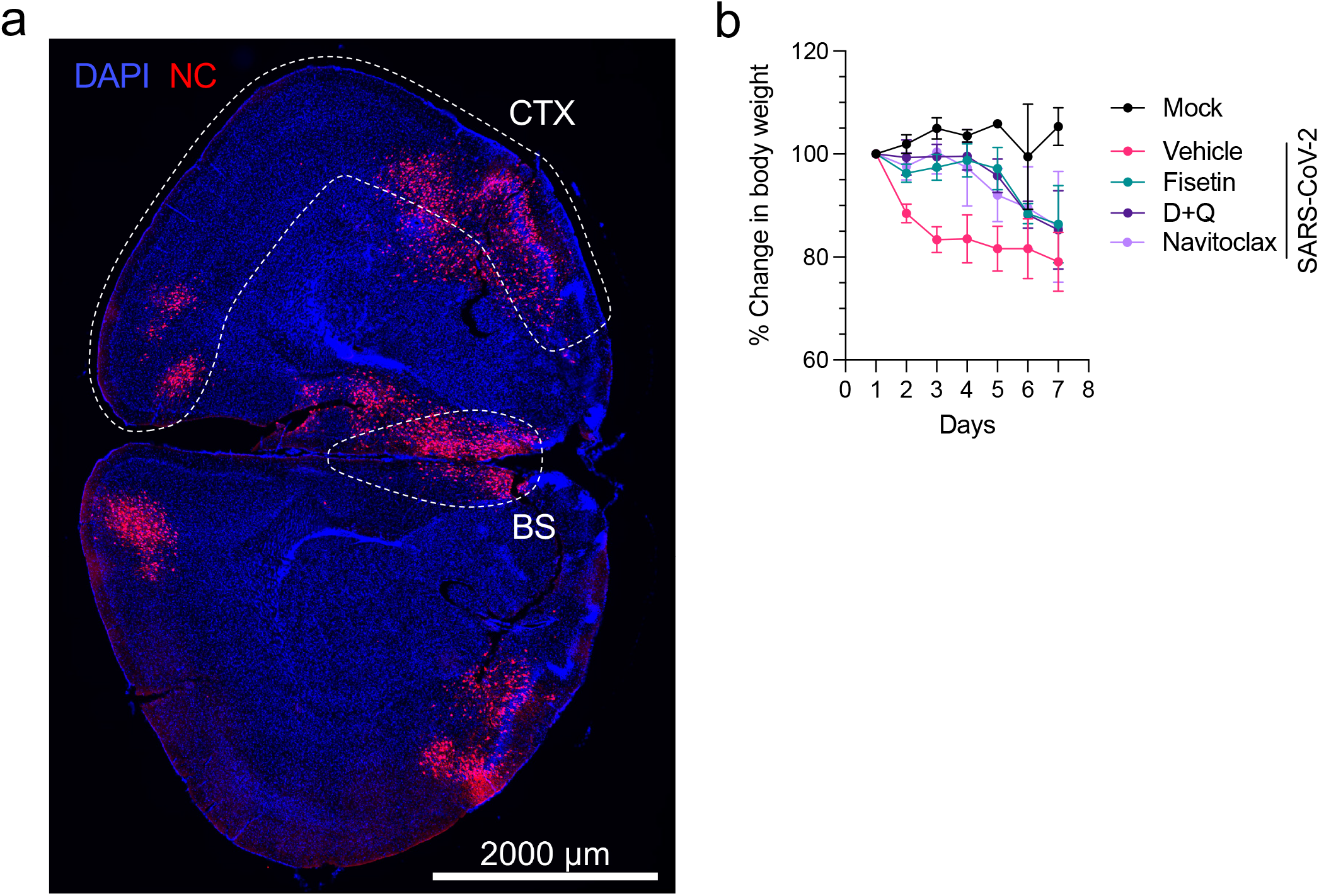
**(a)** Representative immunofluorescent images of viral nucleocapsid (NC) antigen in whole brain coronal sections of brains from SARS-CoV-2-infected K18-hACE2 transgenic mice (5 days post infection). CTX: Cerebral cortex; BS: Brainstem. **(b)** Percentage weight loss up to 7 days post infection. Uninfected mice (n = 3), and Delta SARS-CoV-2-infected mice treated with vehicle (n = 6), fisetin (n = 8), D+Q (n = 8), or navitoclax (n = 8).

**Supplementary Table 1.**
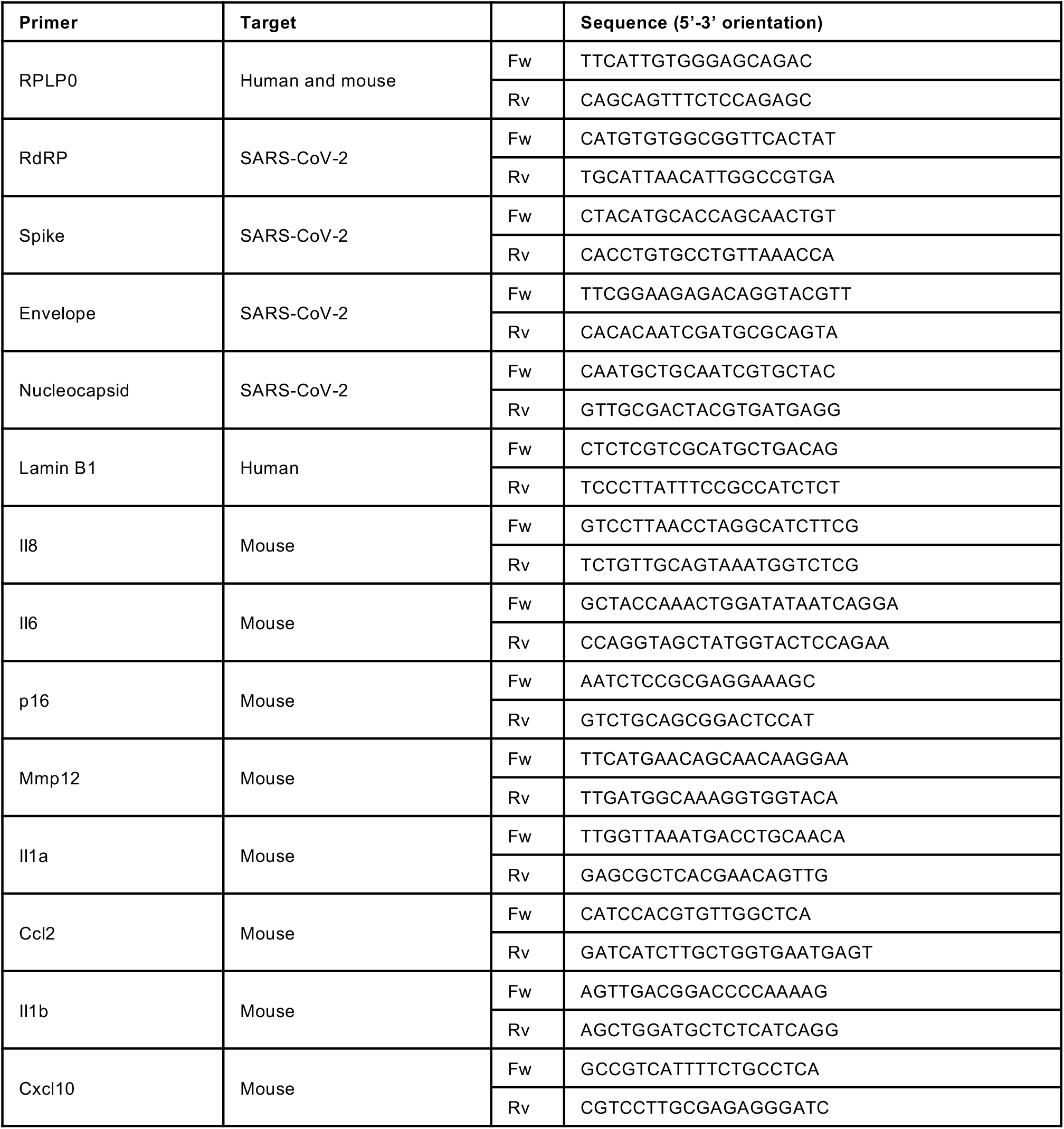

